# Phase-separation of EML4-ALK variant 3 is dependent upon an active ALK conformation

**DOI:** 10.1101/2020.06.11.144485

**Authors:** Josephina Sampson, Mark W. Richards, Jene Choi, Andrew M. Fry, Richard Bayliss

**Affiliations:** School of Molecular and Cellular Biology, Astbury Centre for Structural Molecular Biology, Faculty of Biological Sciences, University of Leeds, Leeds LS2 9JT, UK; Department of Pathology, Asan Medical Center, University of Ulsan College of Medicine, Seoul, Korea; Department of Molecular and Cell Biology, University of Leicester, Lancaster Road, Leicester LE1 9HN, UK

**Keywords:** EML4-ALK, tyrosine kinase inhibitors, phase separation, NSCLC, cancer

## Abstract

Oncogenic fusions involving tyrosine kinases are common drivers of non-small cell lung cancer (NSCLC). There are at least 15 different variants of the EML4-ALK fusion, all of which have a similar portion of ALK that includes the kinase domain, but different portions of EML4. Targeted treatment with ALK tyrosine kinase inhibitors (TKIs) has proven effective but patient outcomes are variable. Here, we focus on one common variant, EML4-ALK V3, which drives an aggressive form of the disease. EML4-ALK V3 protein forms cytoplasmic liquid droplets that contain the signalling proteins GRB2 and SOS1. The TKIs ceritinib and lorlatinib dissolve these droplets and the EML4-ALK V3 protein re-localises to microtubules, an effect recapitulated by an inactivating mutation in the ALK catalytic site. Mutations that promote a constitutively active ALK stabilise the liquid droplets even in the presence of TKIs, indicating that droplets do not depend on kinase activity *per se*. Uniquely, the TKI alectinib promotes droplet formation of both the wild-type and catalytically inactive EML4-ALK V3 mutant, but not in a mutant that disrupts a hallmark of the kinase activity, the Lys-Glu salt-bridge. We propose that EML4-ALK V3 liquid droplet formation occurs through transient dimerization of the ALK kinase domain in its active conformation in the context of stable EML4-ALK trimers. Our results provide insights into the relationship between ALK activity, conformational state and the sub-cellular localisation of EML4-ALK V3 protein, and reveal the different effects of structurally divergent ALK TKIs on these properties.

## INTRODUCTION

Lung cancer is the most common cancer globally and, despite advances in targeted treatment, the 10 year survival rate of patients with lung cancer is only 9% in the UK (Sabir et al., 2017). More than 80% of lung cancers are classified as non-small cell lung cancer (NSCLC), of which most are adenocarcinomas. While activating mutations in RAS or epidermal growth factor receptor (EGFR) are the most common drivers of NSCLC, oncogenic fusions created by chromosome translocations are also frequent driver events. These encode proteins in which the kinase domain of a receptor tyrosine kinase (RTK) is constitutively activated by translational fusion to a self-associating region of another protein. One common example is the fusion of anaplastic lymphoma kinase (ALK) and echinoderm microtubule-associated protein like 4 (EML4), found in ∼2-9% of NSCLC patients (Soda et al., 2007). To date, at least 15 EML4-ALK variants have been identified all containing the same region of ALK. The variants differ in the point of fusion within the EML4 gene but all retain the region of EML4 encoding its N-terminal trimeric coiled-coil domain (TD) through which they achieve self-association and constitutive ALK activation (Soda et al., 2007, Richards et al., 2014). The predominant EML4-ALK variants identified in patients are variants 1 and 3 (V1 and V3), accounting for around 33% and 29% of cases, respectively. Structurally, EML4-ALK V3 contains the TD and an unstructured, basic region that mediates microtubule association of EML4, but lacks its entire C-terminal tandem atypical β-propeller (TAPE) domain, whereas variant 1 also retains a large part of the TAPE domain (Richards et al., 2014).

Since EML4-ALK fusions were identified in NSCLC, potent tyrosine kinase inhibitors (TKI) against ALK have been developed and used as a first-line treatment for those patients. First- and second-generation inhibitors, crizotinib, ceritinib and alectinib are currently used as first-line treatment for ALK-positive NSCLC patients, while lorlatinib, a third-generation inhibitor, is approved for a second or third-line treatment after crizotinib, ceritinib or alectinib in ALK-positive metastatic NSCLC (Solomon et al., 2018, Solomon et al., 2014). NSCLC harbouring different EML4-ALK variants exhibit different responses to ALK inhibitors. NSCLC patients identified with EML4-ALK V1 respond to crizotinib and have increased progression-free survival (PFS) after treatment (Yoshida et al., 2016, Shaw et al., 2011). However, EML4-ALK V3 positive NSCLC patients demonstrate a higher metastatic spread, failure of treatment with ALK inhibitors and increased aggressiveness of the disease, while *in vitro* NSCLC cells harbouring EML4-ALK V3 exhibit resistance to various ALK inhibitors (Christopoulos et al., 2018, O’Regan et al., 2020, Woo et al., 2017).

*ALK* mutations have also been identified in neuroblastoma where, rather than being involved in gene fusions, *ALK* is amplified or carries point mutations within the region encoding the kinase domain (Umapathy et al., 2019). The cancer-related point mutations, most commonly substitutions at F1174 or R1275, are thought to promote an active kinase domain conformation in monomeric ALK (Holla et al., 2017). However, these point mutations alone may be insufficient to drive neuroblastoma, and additional lesions such as *MYCN* amplification may be required (De Brouwer et al., 2010, Jiang et al., 2018). Several of the same mutations confer resistance to TKIs such as crizotinib and frequently arise in the context of ALK fusions in response to TKI treatment (Umapathy et al., 2019).

The cytoplasm is no longer considered a homogenous solution but rather to contain unevenly distributed protein and RNA molecules forming dynamic assemblies through transient molecular interactions. This behaviour sometimes causes molecules associated with these dynamic assemblies to partition from the bulk cytoplasm into droplets through a process called phase separation (Boeynaems et al., 2018, Hyman et al., 2014). Several studies have reported proteins forming droplets within the cytoplasm, such as Tau and polo-like kinase 4 (PLK4) (Wegmann et al., 2018, Park et al., 2019). Liquid-like droplets have also been described in numerous papers as membraneless organelles, such as the P granules in *C. elegans* embryos (Brangwynne et al., 2009). The major characteristics of liquid-like droplets are that they: a) fuse after touching and revert into a spherical shape, b) deform, diffuse and exchange material with the cytoplasm, c) adopt a spherical shape that is driven by surface tension, d) recover rapidly through internal rearrangement when photobleached (Hyman et al., 2014). Recently, Tulpule and colleagues have observed RTK fusion proteins, including EML4-ALK V1 and V3, forming membraneless cytoplasmic granules that act as centres for the organization and activation of RAS and other downstream signalling pathway components (Tulpule et al., 2020)(Preprint). Intriguingly, the granules are disrupted by crizotinib, a first-generation inhibitor of ALK, leading to the hypothesis that their formation is dependent on ALK activity.

In the present study, we demonstrate that EML4-ALK V3 partitions into liquid droplets in the cytoplasm in a manner dependent on the active conformation, rather than catalytic activity, of the ALK kinase domain. We confirm that the droplets capture signalling proteins such as GRB2 and SOS1, and thus may act as a centre for downstream signalling pathways. We investigate how the ALK inhibitors, ceritinib, alectinib and lorlatinib, differentially affect the localisation and behaviour of EML4-ALK V3 inside the cell by dissolving or maintaining these liquid droplets, and describe structure-function relationships between mutated forms of ALK and formation of the liquid droplets.

## RESULTS

### Active EML4-ALK V3 forms cytoplasmic liquid droplets

Previous studies have reported distinct subcellular localisation of EML4-ALK to cytoplasm or microtubules depending on the type of variant (Richards et al., 2015, Hrustanovic et al., 2015). Here, the sub-cellular localisation of EML4-ALK V3 was examined by fixed-cell imaging in several cell lines: a NSCLC patient-derived cancer cell line harbouring endogenous EML4-ALK V3 (H2228), a non-transformed human lung epithelial cell line (Beas2B) expressing EML4-ALK V3 in a doxycycline-inducible manner and HEK293 cells overexpressing YFP-EML4-ALK V3. We identified 25-40 droplet-like structures of concentrated EML4-ALK V3 in the cytoplasm of each of these three different cell types (Fig. 1A, B). However, the YFP-EML4-ALK V3 kinase dead D1270N mutant (KD) exhibited more prominent microtubule binding and fewer droplets (mean of 12 per cell) in transfected HEK293 cells (Fig. 1A, B). In contrast, YFP-ALK 1058-1620 (the region of ALK present in the EML4-ALK fusions) showed very few (at most 5) droplets in the cytoplasm upon expression in HEK293 cells (Fig. S1A, B). Taken together, these observations suggest that an active ALK kinase domain and the EML4 fusion partner are both required for liquid droplet formation.

**Figure 1.**
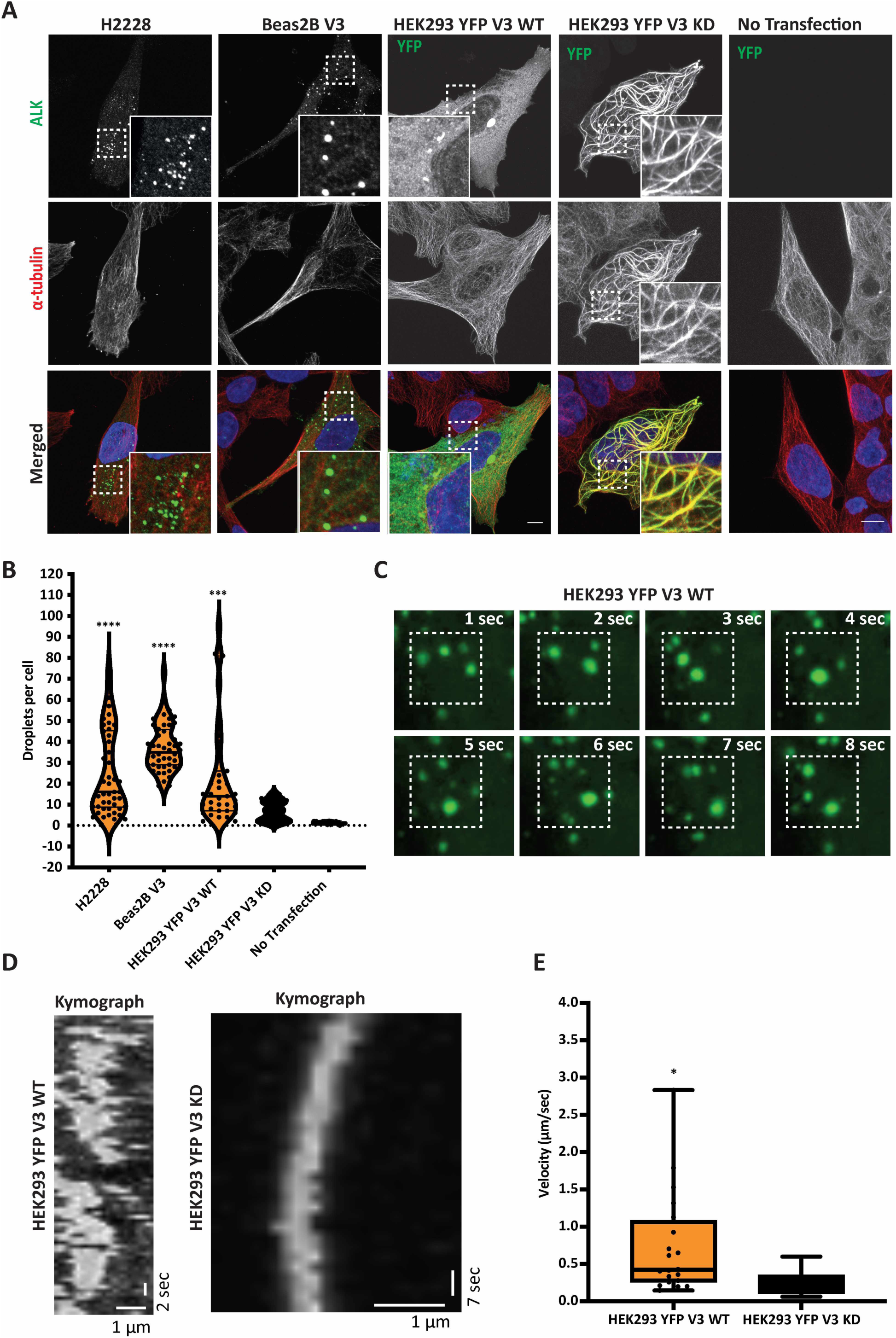
EML4-ALK V3 activation induces formation of cytoplasmic droplets. **A.** H2228 cells expressing endogenous EML4-ALK V3, Beas2B cells stably expressing inducible EML4-ALK V3, and HEK293 transfected with YFP-EML4-ALK V3 WT and KD were stained for either anti-ALK or anti-GFP (green), anti-α-tubulin (red), and DAPI (blue). **B.** Violin plot showing the number of EML4-ALK V3 droplets per cell. \*\*\**P*<0.001, \*\*\*\**P*<0.0001 in comparison to HEK293 YFP-EML4-ALK-V3 KD by one-way ANOVA analysis. **C.** Time-lapse imaging of transfected HEK293 YFP-EML4-ALK V3 WT to observe the movement of droplets. Representative still images are shown of an area of cytoplasm containing YFP-EML4-ALK V3 WT droplets at the times indicated. Scale bar, 10 μm. **D.** Kymographs showing the movement of YFP-EML4-ALK V3 WT droplets and microtubules decorated with YFP-EML4-ALK V3 KD over the duration of the observation period. Scale bars; 1 μm horizontal; 2 or 7 seconds vertical. E. Whisker plot showing the calculated velocity of single events in the kymograph. Data represent 20 counts from 10 kymographs. *n*= 2. \**P*< 0.05 in comparison to DMSO by unpaired *t* test.

In fixed cells, EML4-ALK V3 droplets have a smooth, round appearance as would be expected for liquid structures. To observe whether they also exhibit the accepted dynamic properties of liquid droplets, the movements of individual EML4-ALK V3 droplets in live cells were monitored by time-lapse imaging. YFP-EML4-ALK V3 WT droplets were observed to split and coalesce frequently and to diffuse in a random manner in the cytoplasm with maximum speeds in the range 0.5-3 μm/sec, inversely correlating with droplet size (Fig. S1C-F). While live cell imaging confirmed the presence of distinct cytoplasmic droplets in HEK293 cells transfected with YFP-EML4-ALK V3 WT (Fig 1C), these were not present with the KD construct (Supplementary Movie 1 and 2). We generated kymographs to represent the movements of YFP-labelled structures in live cells revealing that YFP-EML4-ALK V3 WT droplets moved erratically at a mean velocity of 0.7 μm/sec, while microtubules decorated with YFP-EML4-ALK V3 KD drifted smoothly at a mean velocity of 0.2 μm/sec (Fig. 1D-E). These observations suggest that the dynamic cytoplasmic droplets formed by EML4-ALK V3 are a form of liquid phase separation rather than insoluble protein aggregates.

Because kinase dead ALK displayed much reduced phase separation, and activation of RTKs proceeds through dimerization of the kinase domain (Lemmon and Schlessinger, 2010), we hypothesized that direct dimerization of active ALK kinase domains in the context of trimeric EML4-ALK V3 may drive liquid droplet formation. To test this concept we took advantage of a chemical dimerization system using Coumermycin A1 to induce dimerization of GyrB-fused proteins, which has been used to study the dimerization/activation of signalling receptors such as RAF or JAK2 (Farrar et al., 1996, Mohi et al., 1998). A YFP-tagged construct was made in which residues 1-222 of EML4, which is the portion of EML4 present in V3, was fused to GyrB rather than to the ALK kinase domain (Fig. 2A). Hence, in this case dimerization of GyrB by Coumermycin A1 reproduces the effect of active ALK kinase domain dimerization. In untreated HEK293 cells this protein was bound to microtubules with excess material being diffusely distributed in the cytoplasm, but in the presence of Coumermycin A1 YFP-EML4-GyrB was observed to condense into cytoplasmic droplets, at an average of 117 droplets per cell (Fig. 2B, C). These data suggest that liquid droplet formation is dependent upon dimerization of protomers that are already bundled into trimers through the EML4 TD and arises from this stoichiometric mismatch. We therefore propose a model in which EML4-ALK V3 protomers are able to organise themselves, through stable trimerization via the EML4 TD and transient dimerization via the ALK kinase domain in its active conformation, into a dynamic molecular network that partitions from the bulk cytoplasm into distinct cytoplasmic droplets (Fig. 2D).

**Figure 2.**
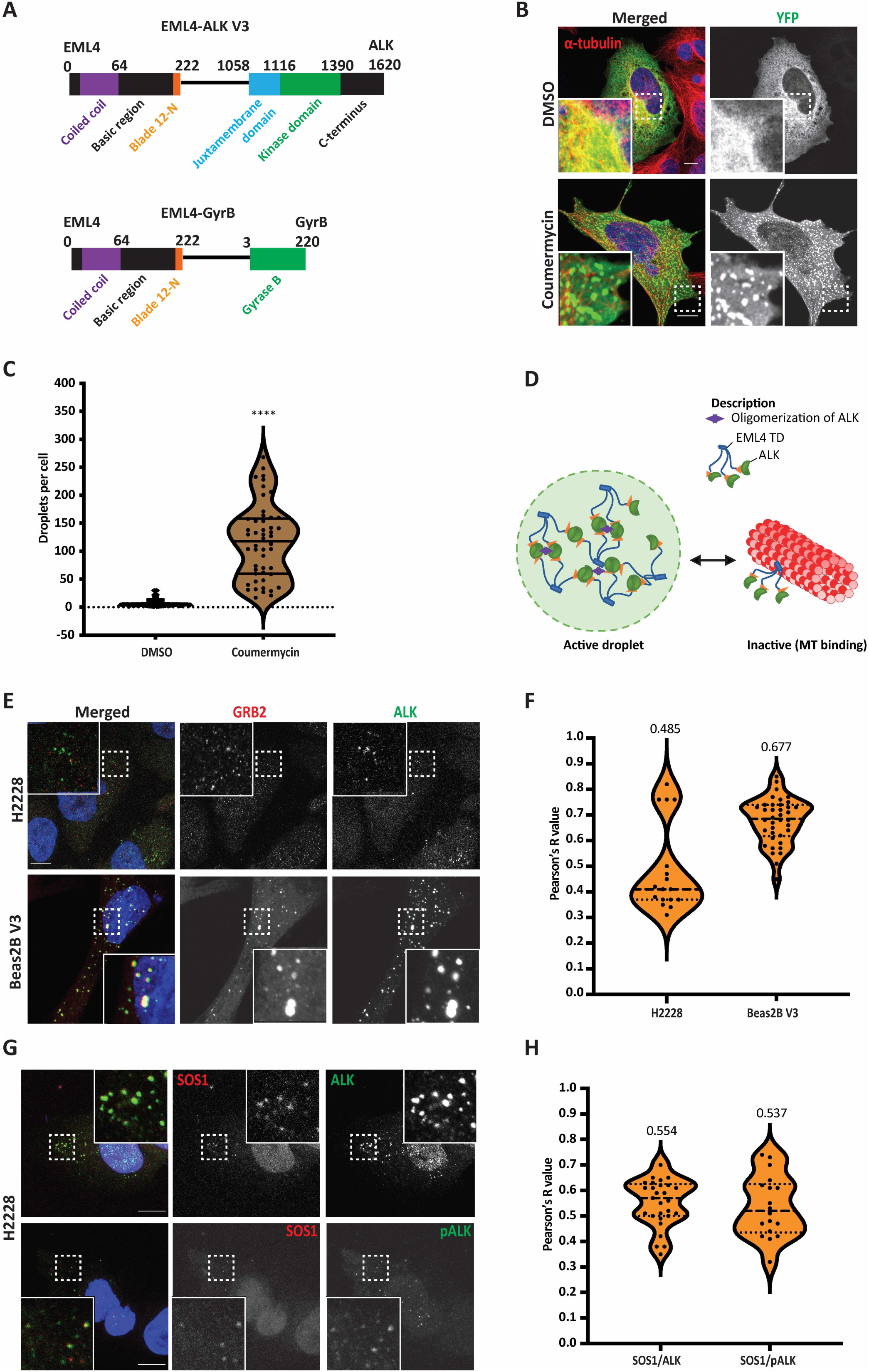
EML4-ALK V3 droplets contain downstream signalling proteins. **A.** Linear representation of the structure of EML4-ALK V3 and EML4-GyrB fusion proteins including residue numbering indicating domain boundaries. **B.** HEK293 cells transfected with YFP-EML4-GyrB and either untreated (DMSO) or treated with Coumermycin A1 to induce dimerization of GyrB before fixation and staining as in A. Scale bars, 10 μm; magnified views of a selected area are shown. **C.** Violin plot showing the number of droplets per HEK293 cell transfected with YFP-EML4-GyrB. Data represent counts from >30 cells, *n*= 4. \*\*\*\**P*< 0.0001 in comparison to DMSO by unpaired *t* test. **D.** A working model for the behaviour of EML4-ALK V3 in which dynamic cross-linking of EML4-ALK trimers arises from the stoichiometric mismatch between the trimeric EML4 N-terminus and the dimers formed by ALK in its active conformation which leads to the formation of liquid droplets in the cytoplasm that exclude large complexes such as microtubules. When the ALK kinase domain is monomeric cross-linking does not occur and EML4-ALK V3 trimers instead localise to microtubules through the EML4 TD and basic region. **E.** H2228 and **G.** Inducible Beas2B V3 cells were stained for anti-ALK or anti-pALK (green), anti-GRB2 or anti-SOS1 (red) and DAPI (blue). **F, H.** Intensity profiles showing co-localisation between GRB2 or SOS1 and ALK or pALK staining. *R* (Pearson’s correlation coefficient) measures the correlation between GRB2 or SOS1 and ALK or pALK signals. Pearson’s measurements from 20-30 droplets of 10 cells for each antibody combination. Data in all violin plots represent counts from at least 20 cells, *n*= 2 or 3. \*\*\*\**P*< 0.0001 in comparison to DMSO by one-way ANOVA.

### EML4-ALK V3 droplets contain RAS/MAPK pathway signalling proteins

We hypothesised that phase separation should not segregate EML4-ALK V3 away from other cytoplasmic components that are required for oncogenic, downstream signalling pathways, and that these must therefore be recruited into the droplets. Although proteomic analyses and cellular assays have identified several cellular networks through which EML4-ALK may signal (Zhang et al., 2016, Hrustanovic et al., 2015), we chose to look for components of the RAS/MAPK signalling pathway that have been located in EML4-ALK V1 and V3 cytoplasmic granules by Tulpule et al. (Tulpule et al., 2020); preprint). Using immunofluorescence microscopy, we compared the localisation of MAPK pathway components, GRB2 and SOS1, with EML4-ALK V3. We observed localisation of endogenous GRB2 to the cytoplasmic EML4-ALK V3 droplets, although co-localisation was less pronounced in the patient-derived H2228 cells than in isogenic Beas2B cells that inducibly express EML4-ALK V3 (Fig. 2E, F). Endogenous SOS1 protein also localised to EML4-ALK V3 droplets in H2228 cells, showing co-localisation both with total ALK and active ALK, phosphorylated on Tyr1604 (Fig. 2G, H). We next used a proximity ligation assay (PLA) to visualise incidences of pair-wise interactions between EML4-ALK V3 and GRB2 in the cytoplasm of Beas2B cells and observed that the occurrence of these foci was significantly reduced in the presence of ALK inhibitors, ceritinib, alectinib and lorlatinib, as would be expected for a signalling complex mediated by SH2 domain/phosphopeptide interactions (Fig. S2). Together, these results show clear recruitment of MAPK pathway components to the EML4-ALK V3 droplets supporting the model that these droplets act as signalling hubs in the cytoplasm.

### Different ALK inhibitors promote either microtubule binding or liquid droplets of EML4-ALK V3

While targeting the EML4-ALK fusion in NSCLC patients with the TKIs, crizotinib, ceritinib and alectinib, has been broadly successful, cancers harbouring EML4-ALK V3 tend to respond less successfully to ALK inhibitors (Kwak et al., 2010, Shaw et al., 2011, Solomon et al., 2014, Woo et al., 2017, Christopoulos et al., 2018). We therefore examined the cellular responses of EML4-ALK V3 to ALK inhibitors. Consistent with the observation that inactive ALK does not form liquid droplets, ALK inhibitors, ceritinib and lorlatinib, dissolved the EML4-ALK V3 liquid droplets and EML4-ALK became more prominently associated with microtubules (Fig. 3A, C). However, alectinib treatment unexpectedly supported the formation of liquid droplets (Fig. 3A, C). Furthermore, alectinib also promoted the formation of liquid droplets with the kinase dead YFP-EML4-ALK V3, which localises to microtubules in the absence of ALK inhibitors or in the presence of ceritinib or lorlatinib (Fig. 3B, D). Quantification of the co-localisation of YFP-EML4-ALK V3 with microtubules confirmed that ceritinib and lorlatinib treatment promotes microtubule localisation of WT or kinase-dead ALK fusion protein (Fig. 3E). In contrast, the co-localisation of EML4-ALK V3 droplets with microtubules in alectinib treated cells was low and similar to the WT protein in untreated cells (Fig. 3E). Consistent with these findings, live cell imaging of HEK293 cells treated with ceritinib or lorlatinib revealed strong microtubule localisation of YFP-EML4-ALK V3, whereas alectinib-treated cells showed a re-organisation of the YFP-EML4-ALK V3 into liquid droplets (Supplementary Movie 3, 4 and 5). In control experiments, YFP-ALK 1058-1620 exhibited neither microtubule binding nor droplet formation and no effect on localisation was observed upon ALK chemical inhibition (Fig. S3A, B, C).

**Figure 3.**
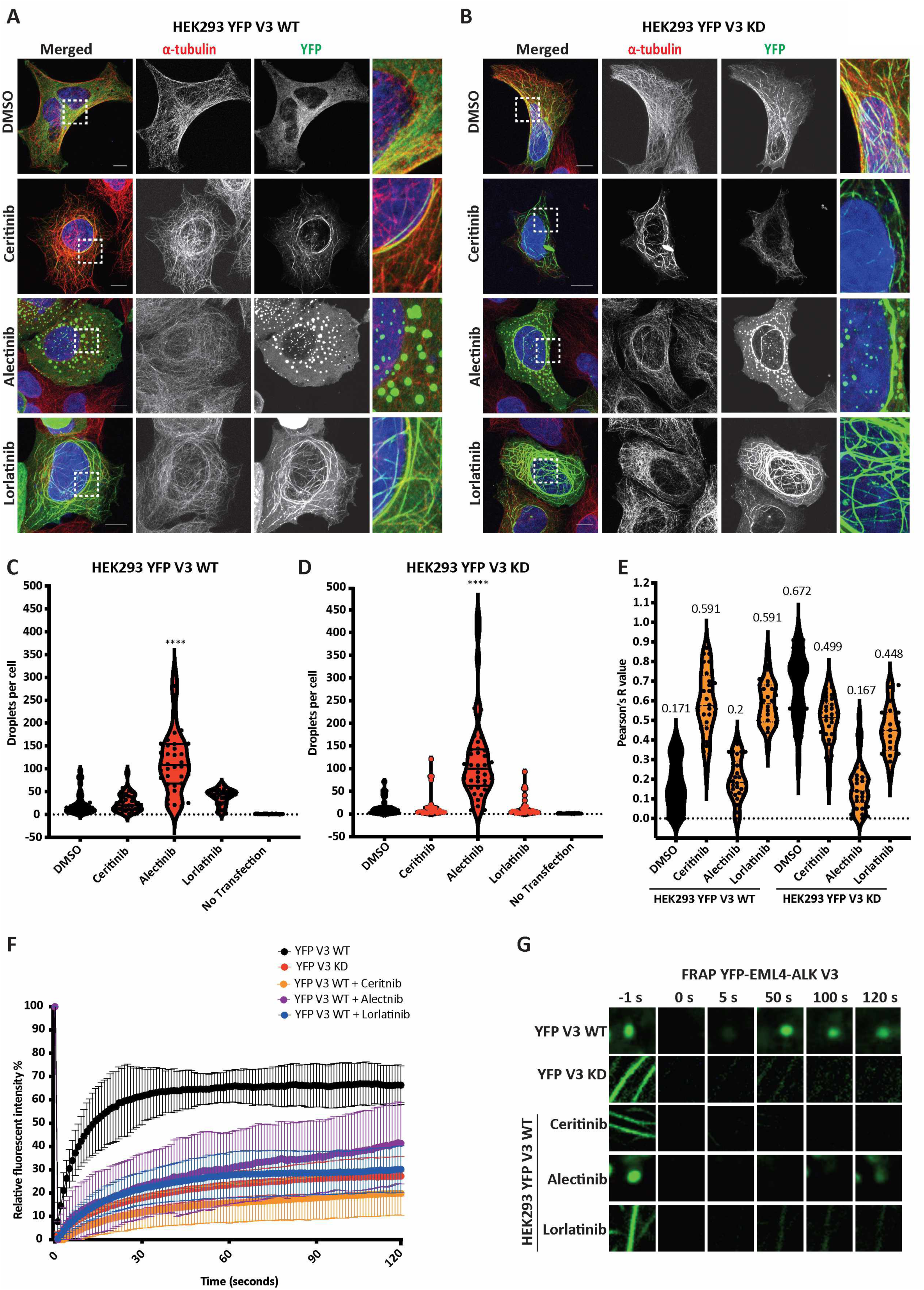
Different ALK inhibitors stabilise either microtubule binding or droplet formation. **A, B.** HEK293 cells were transfected with EML4-ALK V3 WT or Kinase Dead (KD) for 48 hours. Cells were either untreated (DMSO) or treated with ALK inhibitors for 4 hours before fixation and staining with anti-GFP (green), anti-α-tubulin (red), and DAPI (blue). Scale bars, 10 μm; magnified views of a selected area are shown. **C, D.** Violin plots show the number of droplets per cell from A and B, respectively. Data represent counts from >30 cells, *n*= 4. \*\*\*\**P*< 0.0001 in comparison to DMSO by one-way ANOVA. **E.** Intensity profiles showing co-localisation between YFP-EML4-ALK V3 WT or KD and microtubules -/+ ALK inhibitors. *R* (Pearson’s correlation coefficient) measures the correlation between YFP and α-tubulin signals. Pearson’s measurements from 30-50 cells for each construct. **F.** After photobleaching, the fluorescence intensity of an area of a HEK293 cell transfected with YFP-EML4-ALK V3 WT in the presence or absence of ALK inhibitors, or YFP-EML4-ALK V3 KD, was plotted as a function of time. Each curve is the average of data of 40-50 individual cells from 2-3 independent FRAP experiments. **G.** Still images from FRAP analysis of YFP-EML4-ALK V3 WT, KD or ALK inhibitors. Magnified views of a selected area are shown. Time is shown in seconds.

To examine EML4-ALK V3 dynamics within liquid droplets, and its dependence on ALK activity, we used a fluorescence recovery after photobleaching (FRAP) method. The active YFP-EML4-ALK V3 protein (WT, in the absence of inhibitors) recovered rapidly to 70% after bleaching, suggesting that the liquid droplets are highly dynamic structures (Fig. 3F, G) (Supplementary Movie 6). However, inactive YFP-EML4-ALK V3 (WT protein in the presence of ALK inhibitors, ceritinib and lorlatinib, or the mutant protein with the D1270N inactivating mutation) showed very slow recovery on microtubules and also much higher co-localisation with microtubules than active YFP-EML4-ALK V3 (Fig. 3E, F, G). In cells treated with alectinib, YFP-EML4-ALK V3 WT droplets exhibited a faster FRAP recovery than the inactive proteins localised to microtubules, but much slower than the droplets formed by active and uninhibited fusion protein (Fig. 3F, G) (Supplementary Movie 7), and co-localisation of YFP-EML4-ALK V3 WT with microtubules was almost as low in the presence of alectinib as in untreated cells (Fig. 3E). As immunoblotting with phospho-ALK antibodies (pY1604) confirmed that all three inhibitors blocked the kinase activity of EML4-ALK V3 (Fig. S4A, B), the localisation of EML4-ALK V3 to droplets in cells treated with alectinib suggested that ALK catalytic activity *per se* does not drive liquid droplet formation. This led us to investigate the molecular mechanism that may underpin the different responses to these ALK inhibitors.

### ALK maintaining an active kinase conformation is critical for liquid droplet formation

To gain insights into the molecular basis of the different effects of ALK inhibitors, we compared the crystal structures of the unphosphorylated ALK kinase domain in complex with ceritinib (PDB code 4MKC) (Friboulet et al., 2014), lorlatinib (PDB code 4CLI) (Johnson et al., 2014) and alectinib (PDB code 3AOX) (Sakamoto et al., 2011). All three structures show the same overall conformation of ALK, as exemplified by the alectinib-bound structure (Fig. 4A). The ATP binding site, situated between the two lobes of the kinase domain, is occupied by alectinib (coloured magenta in Fig. 4A). The kinase activation loop (coloured blue in Fig. 4A) is partially ordered and adopts a helical conformation that would not be present in the phosphorylated, active kinase. Two other features of the structure, however, are consistent with a kinase in an active state: a set of four hydrophobic residues, termed the regulatory (R-) spine are aligned (coloured green in Fig. 4A), and a salt-bridge between Lys1150 on the β3 strand and Glu1167 on the αC-helix is present (coloured red in Fig. 4A). Close examination of the structures (Fig. 4B, C), and the electron density maps on which these models are based (Fig. 4D, E, F), shows differences in these two features between alectinib and the other two inhibitors. Firstly, one component of the R-spine (Ile1171) and Glu1167 have their side chains in different orientations. Secondly, while the presence of the Lys-Glu salt-bridge in the alectinib structure is well-supported by the experimental data, this feature is much less clear in the other two structures. These differences may reflect the distinct shapes of these inhibitors and their interactions with ALK. Alectinib penetrates deep into the kinase structure within contact range of Ile1171 and its flat shape is compatible with the Lys1150 in a salt-bridge position (Epstein et al., 2012). In contrast, both lorlatinib and ceritinib, and indeed most other ALK TKIs, are non-planar and penetrate less deeply into the kinase structure. This analysis suggests that alectinib promotes an active conformation of the ALK kinase domain while the other inhibitors do not. Based on this analysis, we propose that the ability of alectinib to induce EML4-ALK V3 liquid droplet formation arises from its stabilisation of an ALK conformation characteristic of an active state.

**Figure 4.**
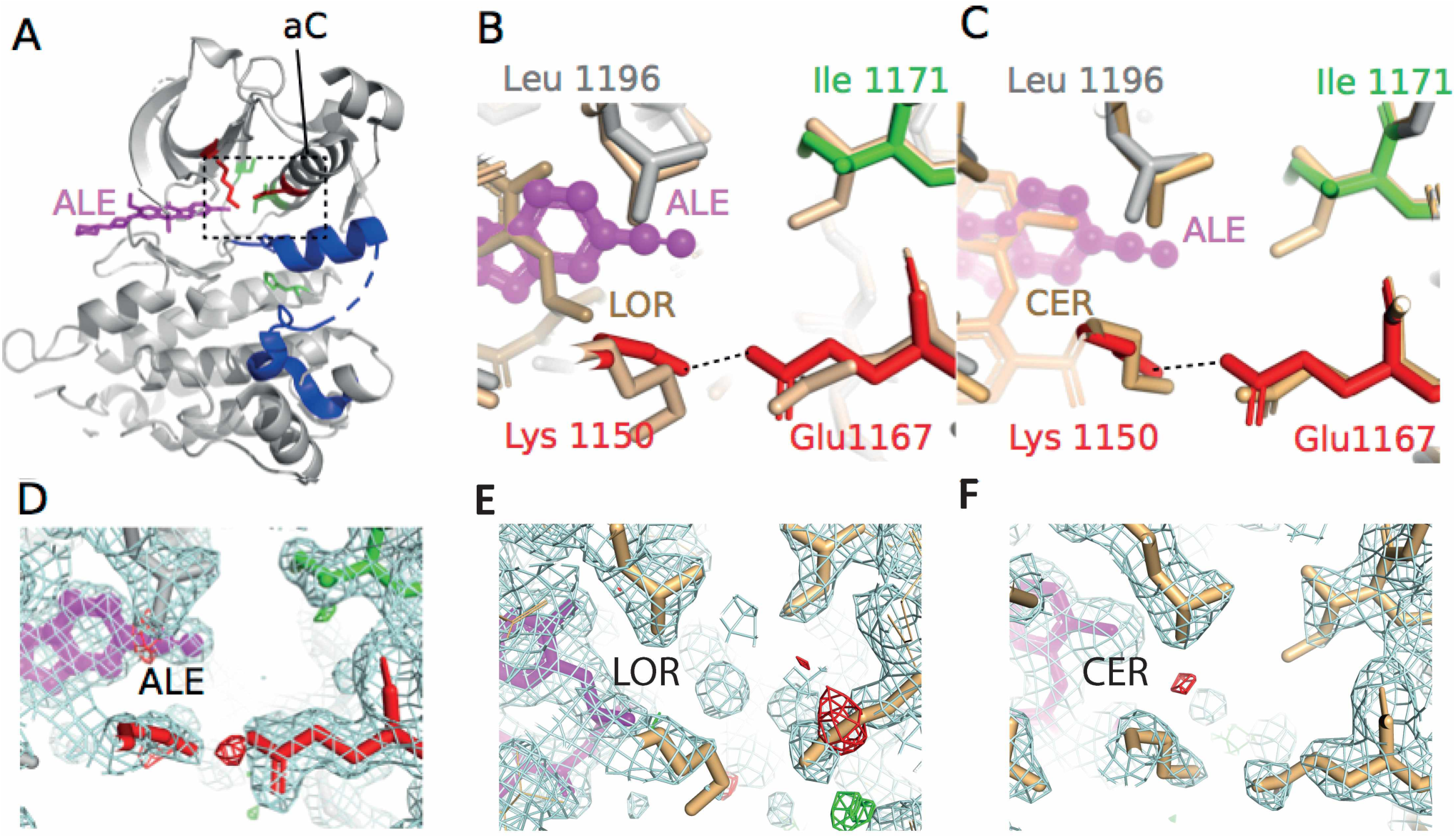
Analysis of ALK structures in the presence of TKIs. **A.** View of alectinib (magenta) bound to ALK (PDB:3AOX) centred on the active site highlighting the αC-helix (blue) and salt bridge (red). **B, C.** Magnified views of active site with key residues shown emphasizing the Lys-Glu salt bridge (red) and alectinib (ALE), lorlatinib (LOR) and ceritinib (CER) binding to it. **D, E, F.** Electron density maps of salt bridge of ALK kinase with alectinib (D), lorlatinib (E), ceritinib (F). 2Fo-Fc maps are coloured light blue and contoured at 1σ. Fo-Fc maps are coloured green (positive) and red (negative) and contoured at ± 3σ. ALK inhibitors are coloured magenta.

To test the hypothesis that an active ALK kinase domain conformation is required for droplet formation, we examined the behaviour of the K1150M mutant incapable of forming the salt-bridge between K1150 and E1167 (coloured red in Fig. 4A), that is a key hallmark of the active conformation of protein kinases. This mutant exhibited strong microtubule binding and produced very few droplets in the presence of all three ALK inhibitors, including alectinib (Fig. 5A, B). The inability of alectinib to rescue the formation of liquid droplets in the salt-bridge mutant, contrasted with the rescue of the active site mutant D1270N, suggested that the salt bridge has a key role in formation of those droplets.

**Figure 5.**
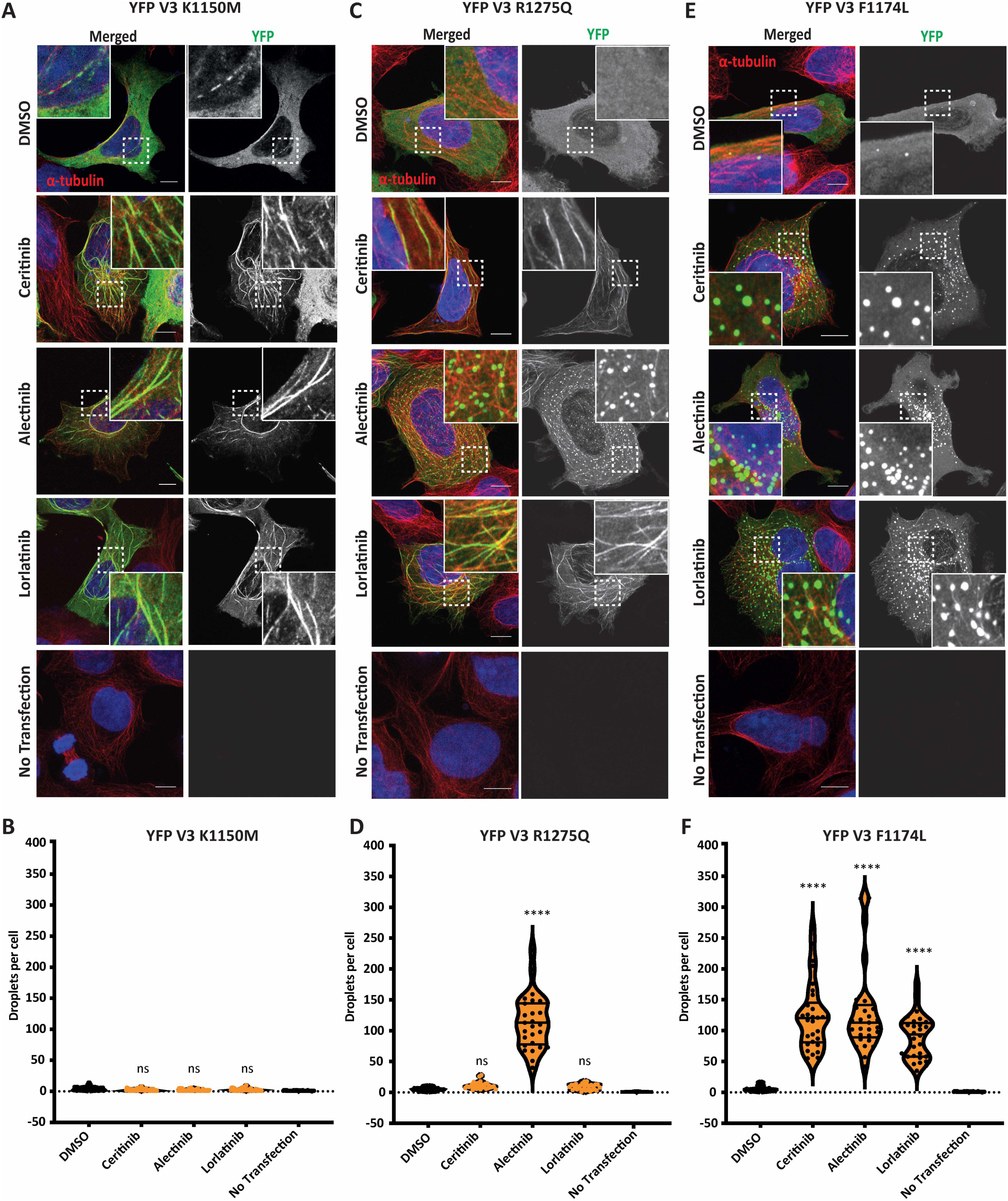
Effect of ALK point mutations on the localisation of EML4-ALK V3. HEK293 cells were transfected with **A.** YFP-EML4-ALK V3 K1150M, **C.** R1275Q or **E.** F1174L constructs and treated with ALK inhibitors or DMSO for 4 hours before fixation and staining with anti-GFP (green), anti-α-tubulin (red), and DAPI (blue). Scale bars, 10 μm; magnified views of a selected area are shown. **B, D, F.** Violin plots representing the number of droplets counted per cell from A, C and E. Data represent counts from at least 30 cells, *n*=3 or 4. \*\*\*\**P*< 0.0001 in comparison to DMSO by one-way ANOVA.

### Somatic ALK mutations associated with neuroblastoma alter the localisation of EML4-ALK V3

The most common *ALK* mutations identified in neuroblastoma occur at Phe1174 (mutated to C, I, S, V, or L) and Arg1275 (L and Q), together representing about 85% of all *ALK* mutations (Holla et al., 2017). To further examine the role of kinase domain conformation in droplet formation, we next investigated whether the F1174L and R1275Q mutations, both of which enhance ALK catalytic activity (Lee et al., 2010, Bossi et al., 2010, Bresler et al., 2014), affect the localisation of the EML4-ALK V3 in the presence of ALK inhibitors. EML4-ALK V3 R1275Q behaved similarly to WT: binding to microtubules in the presence of ceritinib and lorlatinib, and forming droplets (mean of 115 per cell) with alectinib (Fig. 5C, D). YFP-EML4-ALK V3 F1174L exhibited a diffuse cytoplasmic localisation in the DMSO control, however this changed drastically in the presence of ceritinib, alectinib and lorlatinib, which all induced the formation of liquid droplets within the cell (Fig. 5F). More than 90 EML4-ALK V3 F1174L droplets were counted per cell in the presence of all three ALK inhibitors (Fig. 5E, F). Immunoblotting confirmed that all three inhibitors were active against YFP-EML4-ALK V3 R1275Q (Fig. S4C, D), whereas YFP-EML4-ALK F1174L was less sensitive to all three inhibitors (Fig. S4E, F). These results suggest that the mechanism by which the F1174L mutation (but not R1275Q) activates ALK kinase activity involves the promotion of an active conformation of the kinase domain that is competent to drive self-association and droplet formation.

The R1275 residue of ALK (blue, Fig. 6A) is in the activation loop (A-loop; orange, Fig. 6A) and forms interactions with the αC-helix (cyan, Fig. 6A) as observed in crystal structures of ALK in an inactive conformation (Fig. 6A). The F1174 residue of ALK is located on the αC-helix and forms part of a hydrophobic/phenylalanine (Phe) core (black, Fig. 6A) that interacts with an ordered segment of the juxta-membrane (JM) region (green, Fig. 6A) in the crystal structures of ALK in an inactive conformation. We supposed that the F1174L mutation might promote droplet formation through destabilisation of the Phe core and release of the JM segment. To test this hypothesis, we determined the localisation of YFP-EML4-ALK ΔJM (Δ1058-1100). Like the YFP-EML-ALK F1174L mutant, the YFP-EML-ALK ΔJM protein formed droplets (mean of 55 droplets per cell) in the presence of ceritinib that did not co-localise with microtubules (Fig. 6B, C). Hence, we suggest that not only destabilisation of the Phe core, but also release of the JM segment is implicated in the formation of droplets.

**Figure 6.**
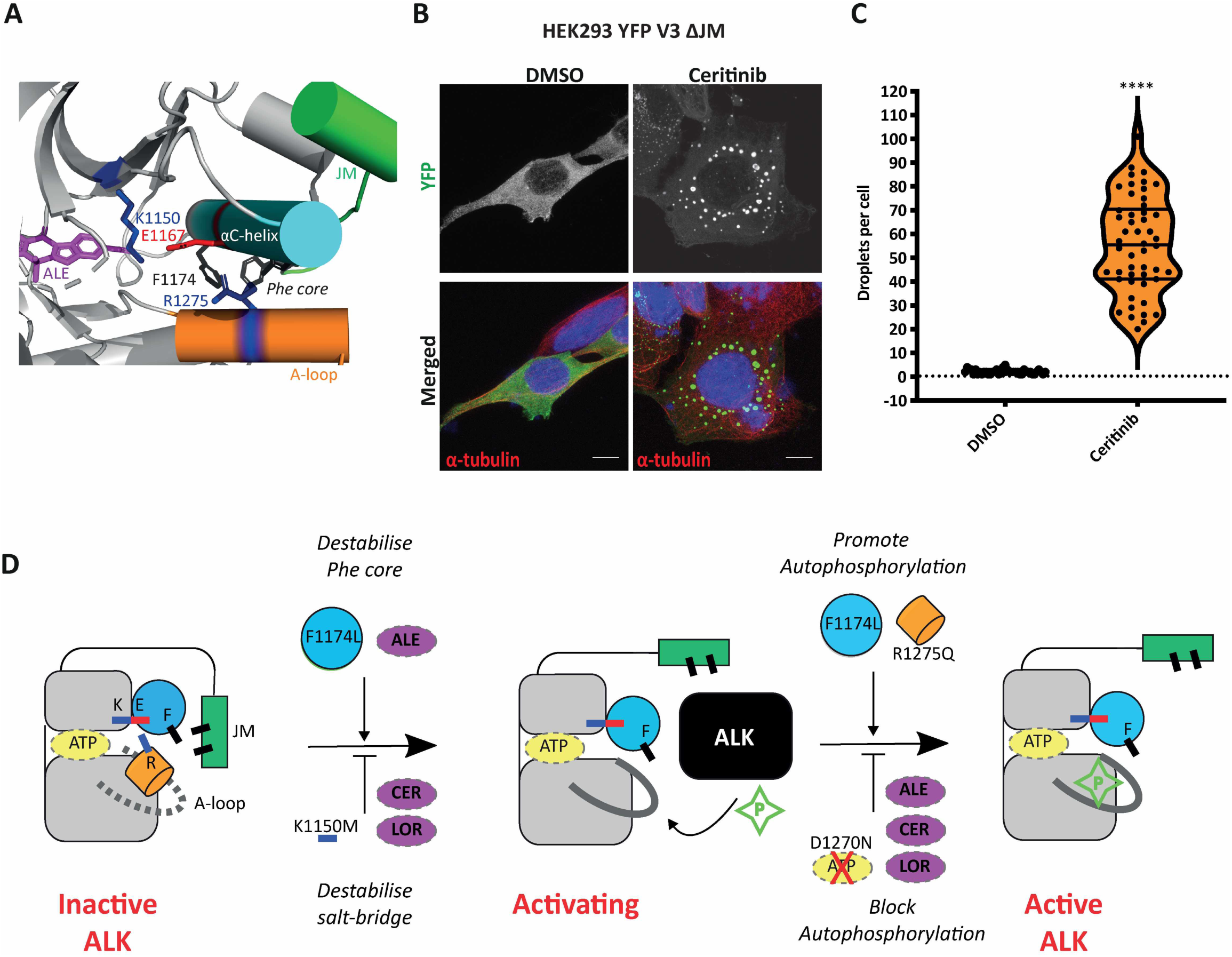
A working model of ALK activation and inhibition. **A**. View of ALK bound to alectinib (magenta), highlighting sites of neuroblastoma mutations and key regulatory features (PDB:3AOX). **B, C.** HEK293 cells were transfected with EML4-ALK V3 ΔJM for 48 hours. Cells were either untreated (DMSO) or treated with ALK inhibitors for 4 hours before fixation and staining with anti-GFP (green), anti-α-tubulin (red), and DAPI (blue). Scale bars, 10 μm; magnified views of a selected area are shown. Violin plots representing the number of droplets counted per cell. D. Cartoon illustrating the ALK kinase with the key residues and domains implicated in autophosphorylation and activation. The salt bridge (blue-red) between K and E residues, the F (cyan) located in the αC-helix and R (orange) located in the activation loop. JM (Juxtamembrane region) is also highlighted. F1147L and alectinib destabilise Phe core, whereas ceritinib, lorlatinib and K1150M destabilise salt-bridge. In addition, F1174L and R1275Q mutations promote autophosphorylation, but ALK inhibitors and D1270N mutation block autophosphorylation.

## DISCUSSION

EML4-ALK V3 in NSCLC is associated with a relatively poor response to ALK inhibitors and more aggressively metastatic disease than other EML4-ALK variants (Christopoulos et al., 2018, Christopoulos et al., 2019, Woo et al., 2017). A better understanding of the molecular biology of EML4-ALK V3 may produce a positive impact for these high-risk NSCLC patients. Here we have demonstrated the ability of EML4-ALK V3 to phase separate to form *de novo* liquid droplets within the cytoplasm and that adoption by ALK of a conformation characteristic of an active state, in which the critical salt bridge between Lys1150 and Glu1167 is intact, is the critical factor for formation of these droplets. The liquid nature of the bodies formed by EML4-ALK V3 is evidenced by their rapid internal rearrangement and their behaviour in the cytoplasm, moving randomly and frequently dividing and coalescing, and stands in contrast to the solid, aggregate-like nature of the particles that Tulpule et al. (2020) have shown are formed by EML4-ALK V1. This is consistent with the key structural difference between EML4-ALK V1 and V3 – the longer variant includes the incomplete globular TAPE domain of EML4 that is predicted to aggregate due to exposure of its hydrophobic core and which confers dependence on the chaperone Hsp90 for stability (Richards et al., 2014).

Analysis of the *de novo* EML4-ALK V3 cytoplasmic droplets reveal that they contain adaptor and signalling proteins, such as GRB2 and SOS1, that function in the RAS/MAPK pathway. We therefore suggest that EML4-ALK V3 achieves downstream signalling within and from cytoplasmic droplets. These findings are consistent with those of Tulpule et al. (Tulpule et al., 2020), who have suggested a similar mode of signalling by which fusions of RTKs, including ALK and RET, direct RAS/MAPK axis signals from membraneless cytoplasmic protein granules. It is notable that more studies now report the existence of cytoplasmic granules or droplets as a mechanism of activation in the cell; for example, the autoactivation of Polo-like kinase 4 by phase separation acts as a mechanism for centriole biogenesis (Park et al., 2019).

We observed striking differences in the localisation of EML4-ALK V3 upon treatment with different ALK inhibitors. We propose that alectinib mimics kinase activation by stabilising the Lys-Glu salt-bridge and an active conformation, promoting phase separation, whereas ceritinib and lorlatinib induce an inactive ALK conformation that disfavours droplet formation and drives microtubule association instead. Stabilisation by alectinib of interactions between ALK kinase domains was also observed in FRAP experiments, where YFP-EML4-ALK V3 in liquid droplets exhibited slower recovery after photobleaching in the presence of alectinib, indicative of reduced dissociation dynamics and mobility. EML4-ALK-positive NSCLC driven by EML4-ALK V3 shows faster metastatic spread than that harbouring other variants (Christopoulos et al., 2018, Woo et al., 2017, O’Regan et al., 2020). This behaviour was recently proposed by O’Regan et al. (2020) to arise from its ability to interact with microtubules, which is not shared by the other variants: EML4-ALK V3 was shown to recruit NEK9 and NEK7 kinases to the microtubule network and to stabilize microtubules, promoting the formation of cellular protrusions and enhancing cell migration. Since this behaviour was not dependent upon ALK activity, chemical inhibition of ALK would not be expected to limit metastasis by this mechanism (O’Regan et al., 2020), and indeed the present work may suggest that those ALK inhibitors, including ceritinib and lorlatinib, that prevent EML4-ALK V3 phase separation might tend to increase metastatic potential in EML4-ALK V3-driven cancer by shifting the protein onto microtubules. However, we propose that patients with cancer driven by EML4-ALK V3 might exhibit a better response with less metastatic spread if treated with alectinib, since it causes the inhibited EML4-ALK V3 protein to be sequestered in cytoplasmic droplets away from the microtubule network.

Crystal structures of ALK reveal the basis for autoinhibition and how it is reversed by activating mutations. In inactive ALK, the JM segment forms a β-turn motif that folds over the kinase domain and interacts with the αC-helix and A-loop via a hydrophobic, phenylalanine (Phe) core that comprises residues such as F1245 and F1174. Mutations of these residues in cancer are strongly activating because they release the autoinhibitory interactions within the Phe core, enabling autophosphorylation to occur (Bresler et al., 2014). A region of the unphosphorylated activation loop (A-loop) of ALK between F1271 and R1279 folds into an α-helix that is incompatible with activating autophosphorylation of Y1278 (Lee et al., 2010, Bossi et al., 2010). Hydrogen bond interactions between the αC-helix and the A-loop helix stabilise the inactive conformation of ALK, and disruption of these interactions through mutations such as R1275Q promotes activation by unwinding the A-loop helix, which releases the side chain of Y1278 (Epstein et al., 2012). We propose a model to summarise our findings and explain how liquid droplet formation and the mechanism of ALK activation through dimerisation of the kinase domain might be related (Fig. 6D). Interactions between ALK kinase domains are favoured by destabilisation of the Phe core by F1174L mutation or alectinib, and disfavoured by disruption of the Lys-Gly salt-bridge by the K1150M mutation or the inhibitors ceritinib and lorlatinib. Productive ALK-ALK interactions result in autophosphorylation and activation within droplets, which may be accelerated through cancer-promoting mutations such as F1174L or R1275Q, while ALK-ALK interactions are non-productive in the presence of alectinib. The structural basis of ALK kinase domain-kinase domain interactions in these droplets is unknown, and requires further study, but we presume that it is related to the mechanism through which ALK and other RTKs are activated.

In a mechanism common among oncogenic RTK fusions, self-association via the EML4 TD at the N-terminus of EML4-ALK V3 mimics the oligomerization-dependent activation mechanism of RTKs and leads to constitutive activation of the ALK kinase domains (Schlessinger, 2000, Lemmon and Schlessinger, 2010). Our model of liquid droplet formation by EML4-ALK V3 extends this by suggesting that the stoichiometric mismatch between the trimeric EML4 portion of the fusion protein and dimers of active ALK kinase domains means that EML4-ALK V3 will tend to cross-link through transient kinase domain interactions when the ALK moiety is in an active conformation, leading to phase separation (Fig. 2H). The droplets are not associated with microtubules, which implies that the interactions between EML4-ALK trimers are incompatible with microtubule binding or that bulky structures such as microtubules are excluded from the liquid droplets. When the ALK moiety is in an inactive conformation, EML4-ALK V3 favours microtubule association through the EML4 moiety, which contains natively microtubule-binding TD and basic regions. Our findings suggest a novel mechanism explaining how the EML4-ALK V3 fusion organises and activates itself to facilitate MAPK downstream signalling, and the importance of the Lys-Glu salt bridge for formation of an active conformation that promotes the formation of cytoplasmic droplets.

## MATERIALS AND METHODS

### Plasmid construction and mutagenesis

EML4-ALK V3 and ALK 1058-1620 were cloned into a version of pcDNA3.1-hygro (Invitrogen) as previously described (Richards et al., 2014), providing an N-terminal YFP tag for transfection of HEK293 cells. D1270N, K1150M, R1275Q and F1174L mutants were generated by QuickChange procedure (Agilent Technologies) and confirmed by sequencing. To generate the ΔJM construct, the region encoding residues 1058-1100 of ALK was deleted from pcDNA3.1-hygro YFP-EML4-ALK-V3 by PCR and the plasmid re-ligated in frame. YFP-EML4-GyrB fusion was constructed by replacing the *ALK* portion of EML4-ALK V3 with a sequence encoding residues 3-220 of GyrB (the ATPase domain of *E. coli* DNA gyrase subunit B).

### Cell culture, transfection and drug treatments

HEK293 (Human embryonic kidney cells), NCI-H2228 (a Human NSCLC cell line harbouring EML4-ALK variant 3b) cells were obtained from ATCC. All cells were obtained within the last 5 years, stored in liquid nitrogen, and maintained in culture at 37°C in a 5% CO_2_ atmosphere for a maximum of 2 months. We relied on the provenance of the original collection for authenticity. Cell lines were regularly tested for mycoplasma contamination using a highly sensitive and specific PCR-based assay (EZ-PCR Mycoplasma kit, Geneflow). NCI-H2228 and Beas2B were cultured in RPMI-1640 medium, and HEK293 in DMEM medium. All media were from Invitrogen-GIBCO and supplemented with 10% heat-inactivated foetal bovine serum (FBS), 100 IU/ml penicillin and 100 mg/ml streptomycin. Beas2B cell line was treated with doxycycline for 72 hours to induce expression of EML4-ALK V3. HEK293 were transiently transfected using Fugene HD (Promega, U.K.) according to manufacturer’s instructions. Ceritinib (LDK378), Alectinib (CH5424802) and Lorlatinib (PF-06463922) were purchased from Selleck Chemicals and stock solutions were prepared in DMSO. Unless otherwise indicated, the following compounds were added to cells for 4 hours: Ceritinib (0.5 µmoles/L); Alectinib (0.1 µmoles /L); Lorlatinib (0.1 µmoles /L); The compounds were diluted in fresh media before each experiment, and control cells were treated with the same volume of DMSO.

### Inducible dimerization of EML4-GyrB fusion

48 hours following transfection with the YFP-EML4-GyrB construct, HEK293 cells were treated for 8 hours with either 0.1 µmoles/L Coumermycin A1 (Promega) or an equivalent volume of DMSO. The average number of droplets formed per cell were measured and analysed by Imaris (v.9.3) software. Data represent the mean of at least three independent experiments ± SD.

### Indirect immunofluorescence microscopy

Cells grown on acid-etched glass coverslips were washed with PBS and fixed with 3.7% formaldehyde in PBS buffer for 10 minutes. Cells were kept in blocking buffer (3% BSA in PBS) for 1 hour and incubated for 2 hours or overnight with primary antibodies diluted in 3% BSA/PBS buffer followed by 1 hour incubation with secondary antibodies. Primary antibodies were against, α-tubulin mouse (1:1000; Sigma), α-tubulin rabbit (1:800; abcam 18251), GFP (1:1000; abcam 6556), ALK rabbit (D5F3) (1:100; CST), ALK (31F12) mouse (1:100; CST), ERK (L34F12) (1:100; CST), pERK (Thr202/Tyr204) (1:100; CST), pALK (Tyr1604) (1:100; CST), GRB2 (Y237) (1:100; abcam 32037), SOS1 (1:100; Santa Cruz Biotechnology). Secondary antibodies were Alexa Fluor-488 and -594 goat anti-rabbit and goat anti-mouse IgGs (1:200; Invitrogen). Imaging was performed on a Zeiss LSM880 + Airyscan Inverted confocal microscope using a 40x oil objective (numerical aperture, 1.4). Z-stacks comprising of 10-20 x 0.3 µm sections were acquired. Images were analyzed using ImageJ (v.2.0.0).

### *In Situ* Proximity Ligation Assay (PLA)

Adherent cells were cultured in the appropriate media and treated with ALK inhibitors for 4 hours before fixation. Cells were fixed with 3.7% (vol/vol) paraformaldehyde for 10 minutes and permeabilized with PBS containing 0.5% Triton X-100 for a further 5 minutes. Cells were incubated with blocking buffer (3% BSA in PBS) for 1 hour and then with the antibodies, as indicated, diluted in the same buffer for overnight. Duolink^®^ proximity ligation assays were carried out according to manufacturer’s instructions (Duolink^®^, Sigma). The average number of PLA dots detected per cell was calculated using DotCount software and data represents the mean of at least two independent experiments ± SD.

### Live cell imaging

Time-lapse imaging was performed using a Zeiss LSM880 + Airyscan Inverted confocal microscope using a 40x oil objective and a scan zoom of 4. Cells were cultured in glass-bottomed culture dishes (Ibidi) and maintained at 37°C in an atmosphere supplemented with 5% CO_2_ using a ZL multi S1 Incubator box system. Drugs were added just before imaging with pre-warmed OptiMEM (ThermoFisher Scientific) containing 10% FBS. Z-stacks comprising of twenty 1 μm steps were acquired every 1 second for 1 minute. Stacks were processed into maximum intensity projections and movies prepared using ImageJ (v.2.0.0).

### Fluorescence Recovery After Photobleaching (FRAP) assay

Live-cell FRAP measurements were made in HEK293 cells expressing fluorescently tagged YFP-EML4-ALK V3 WT or kinase dead (KD) protein. For FRAP, a region of interest (ROI) of 40×40 µm was bleached with 10 iterations at 100% argon 488 nm and 100% 405 nm laser power simultaneously. A second ROI of 40×40 µm was used as control (no photo-bleach) to represent the background. One image was captured prior to bleaching and then images were collected every 1 second for 2 minutes following bleaching. The fluorescence intensity profiles of the ROI were determined using Zeiss software. The corrected fluorescence intensity was calculated by removing the background (no Bleach ROI) using the following equation and was expressed as percentage: (Intensity of ROI – Background Intensity)/Intensity of pre-Bleach

### Colocalization measurements and tracking analysis

Cells stained with GFP/α-tubulin or GRB2/ALK or SOS1/ALK or pALK antibodies for immunofluorescence and 20-30 cells were analyzed for colocalization studies. The colocalization analysis was measured using Colo2 in ImageJ (v.2.0.0) software. A 100 pixels x 100 pixels ROI was positioned over the interphase cell and the Pearson R values were calculated. Data represents the Pearson R values of at least two independent experiments ± SD. For tracking analysis, 12 second movies of HEK293 YFP-EML4-ALK V3 WT droplets were analysed to calculate mean droplet speeds using TrackMate v5.2.0 in ImageJ (Plugins -> Tracking -> TrackMate) (Tinevez et al., 2017). Images were filtered to detect droplets using “LoG detector” (Laplacian of Gaussian) with an estimated blob diameter of 1-2 pixels and a threshold of 1. For kymograph analysis, time-lapse series of YFP fluorescence from HEK293 cells expressing YFP-EML4-ALK V3 WT or YFP-EML4-ALK V3 KD were analysed using ImageJ software. Velocities of individual events were calculated using a custom macro on FiJi. All measurements were taken from at least two independent experiments.

### Cell extracts and Western blotting analysis

Whole-cell lysates were prepared in M-PER mammalian protein extraction reagent (ThermoFisher Scientific) containing a Halt™ protease inhibitor cocktail (ThermoFisher Scientific) and Halt™ phosphatase Inhibitor cocktail (ThermoFisher Scientific). Immunoblot analyses were carried out with anti-phospho-ALK (Tyr1604) (1:1000, 3341; CST) and anti-α-tubulin (1:2000, T5168; Sigma) antibodies. Secondary antibodies were rabbit or mouse horseradish peroxidase -conjugated secondary antibodies (1:1000; Amersham). The blots were visualized using the Pierce™ ECL Western Blotting Substrate (32106; Pierce).

### Statistical analysis

All quantitative data represent means and standard deviation (SD) of at least two independent experiments. Statistical analyses were performed using one-way ANOVA analysis from Prism 7.0 software. ****p<0.0001, ***p<0.001, **p<0.01, *p<0.05.

## ACKNOWLEDGMENTS

We acknowledge support of the University of Leeds Bioimaging and Flow Cytometry facility funded by a Welcome Trust award ‘‘Multifunctional imaging of living cells for biomedical sciences’’. This work was funded by a Programme Award from Cancer Research UK to R.B (C24461/A23302).

## SUPPLEMENTARY MATERIAL

**Supplementary Figure S1.**
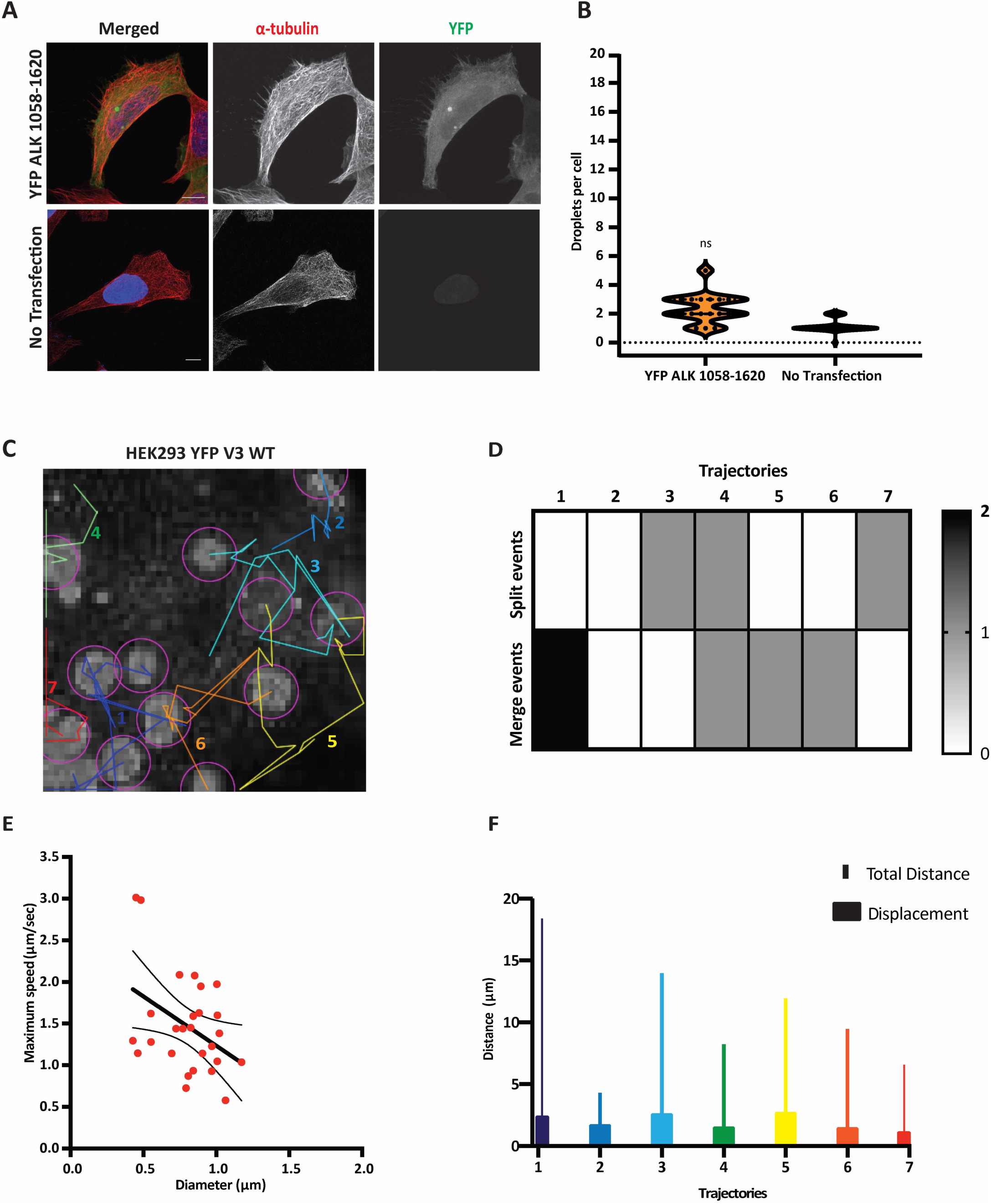
Liquid droplet formation by YFP-EML4-ALK V3 but not YFP-ALK. **A.** HEK293 cells were transfected with YFP-ALK 1058-1620 for 48 hours before fixation and staining with anti-GFP (green), anti-α-tubulin (red) and DAPI (blue). **B.** Violin plots representing the number of liquid droplets per cell. Data represents counts from at least 20 cells, *n*= 2. **C.** Track classification of YFP-EML4-ALK V3 droplets in the cytoplasm of HEK293. Duration of the movie was 12 seconds. Scale bar, 5 μm. Each coloured and numbered trajectory indicates the movement of an individual droplet. **D.** Heatmap representing the number of split and merge events in each trajectory. **E.** Plot of droplet maximum speed (μm/sec) versus droplet diameter (μm). The best-fit line shows correlation with 95% confidence interval. **F.** Plot displays the total distance (μm) and displacement (μm) of each trajectory. Colours and numbers relate to the tracks shown in C.

**Supplementary Figure S2.**
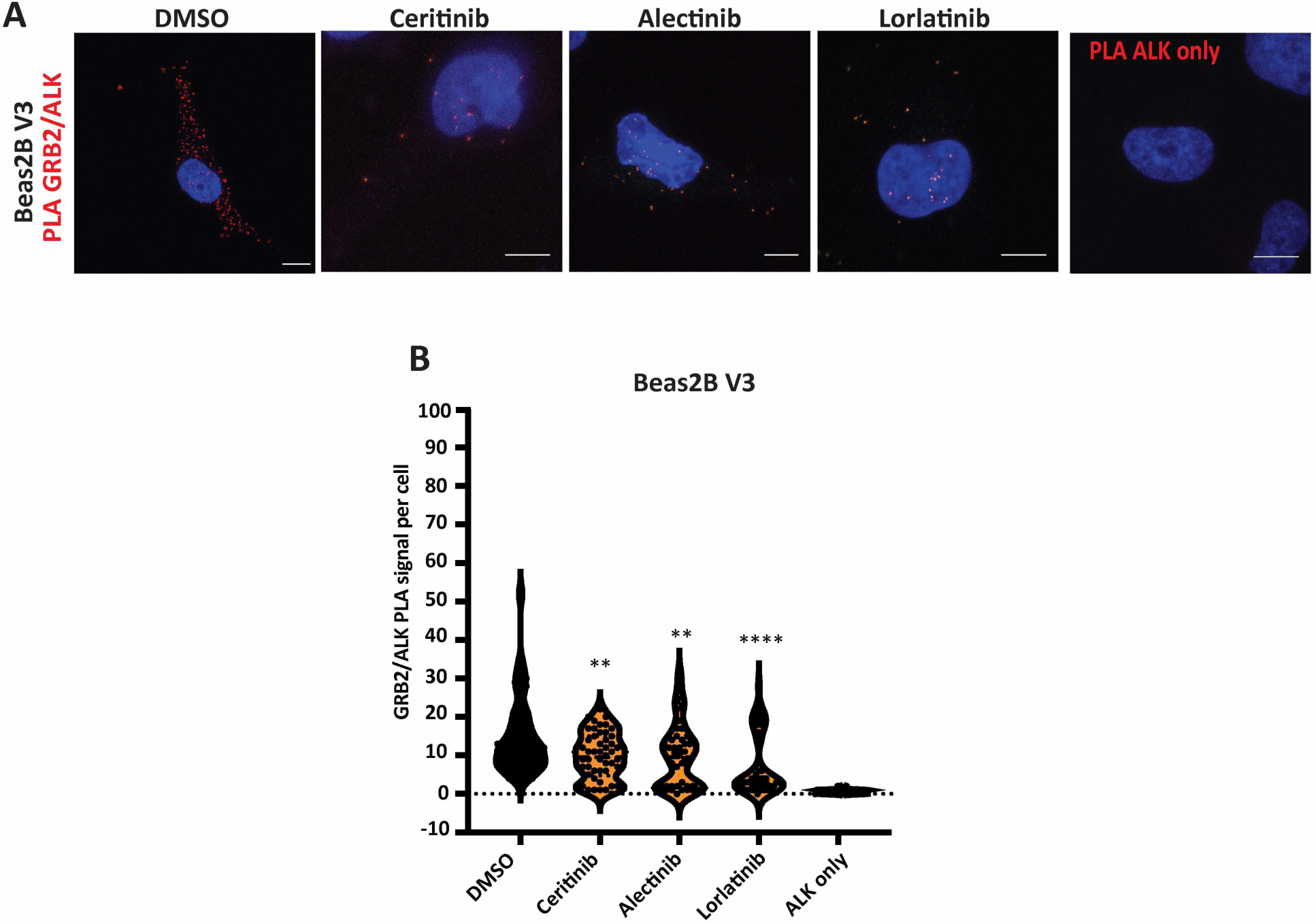
Observation of GRB2/ALK complexes by PLA. **A.** Beas2B V3 cells were treated with ALK inhibitors for 4 hours before fixation and staining with GRB2 and ALK antibodies for PLA. Nuclei are indicated by DAPI staining (blue). Red foci indicate GRB2/ALK protein complexes. Single ALK antibody staining was used as a control for PLA interactions. Scale bars, 10 μm. **B.** Violin plots representing the number of PLA foci per cell from A. Data represent counts from at least 20 cells, *n*= 2.

**Supplementary Figure S3.**
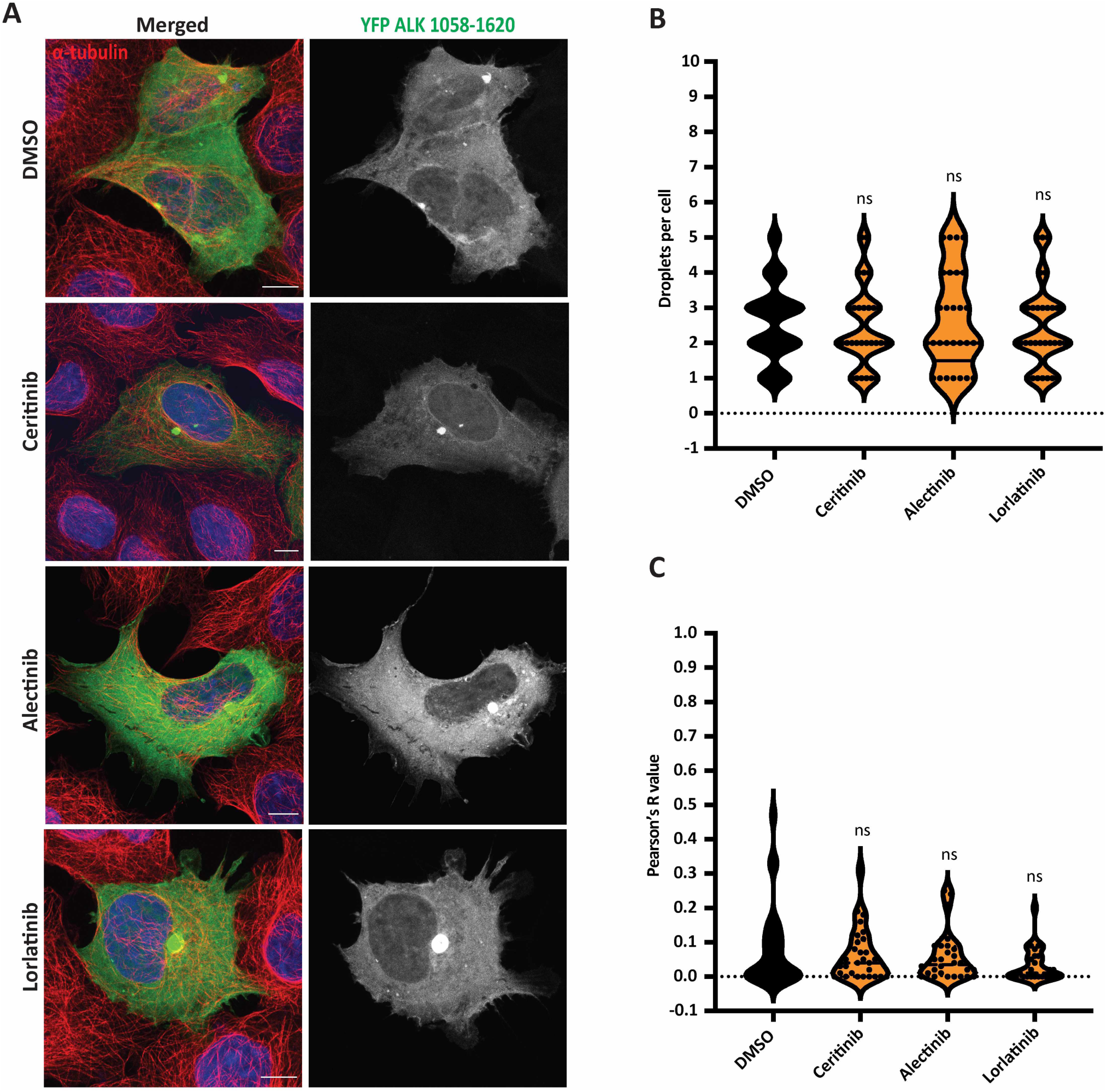
Localisation of YFP-ALK 1058-1620 is unaffected by ALK inhibitors. **A.** HEK293 cells were transfected with YFP-ALK 1058-1620 and treated with ALK inhibitors or DMSO for 4 hours before fixation and staining with anti-GFP (green), anti-α-tubulin (red), and DAPI (blue). Scale bars, 10 μm; magnified views of a selected area are shown. **B.** Violin plot representing the number of droplets per cell from A. Data represent measurements taken from at least 20 cells, *n*= 2. **C.** Violin plot represents intensity profiles showing co-localisation between YFP-ALK 1058-1620 and microtubules in the presence of ALK inhibitors or DMSO. *R* (Pearson’s correlation coefficient) measures the correlation between YFP and α-tubulin signals. Pearson’s measurements from 30-50 cells for each treatment.

**Supplementary Figure S4.**
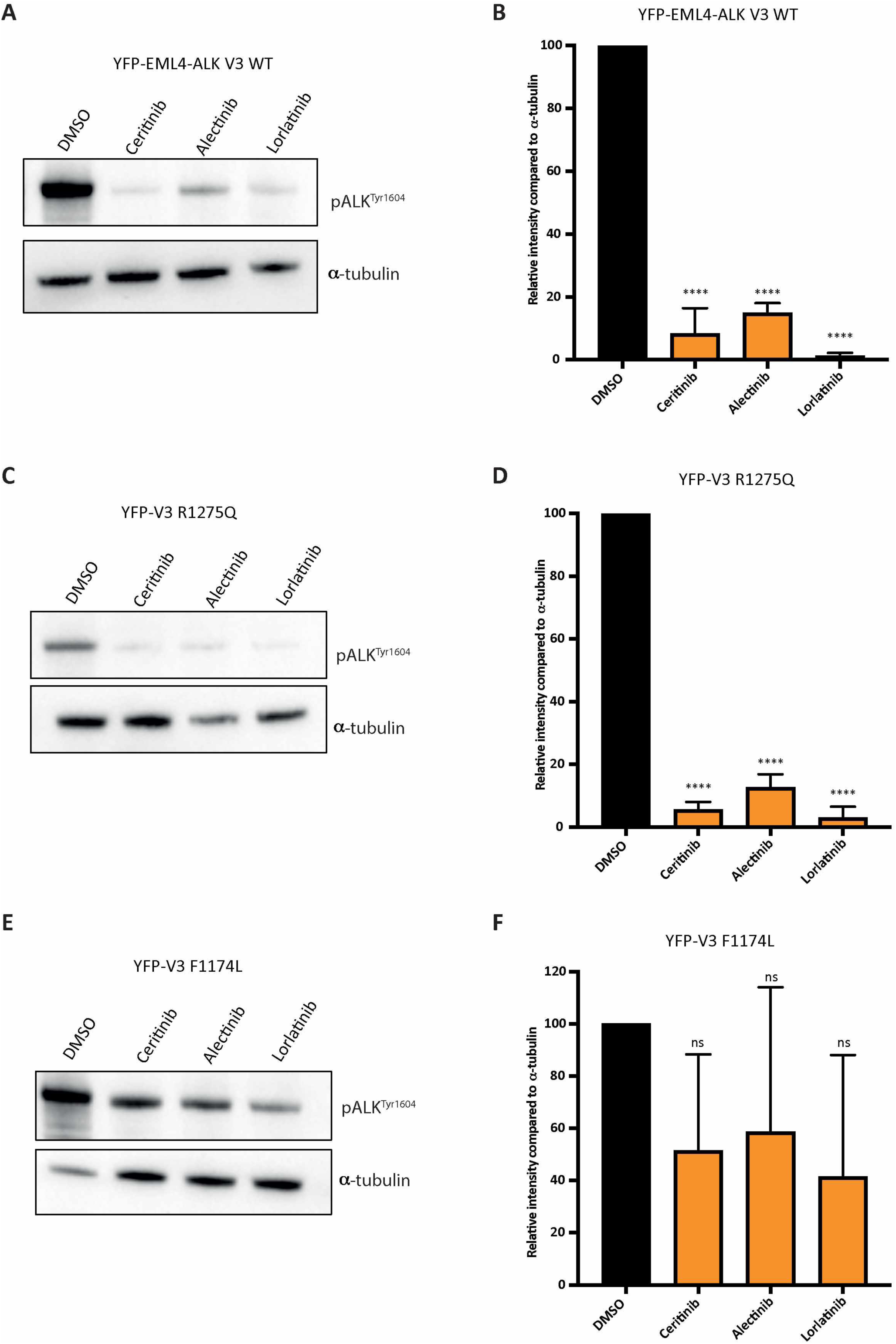
Effect of ALK inhibitors on phosphorylation of EML4-ALK V3 WT and mutants. **A, C, E.** HEK293 cells were transfected with YFP-EML4-ALK V3 WT, V3 R1275Q or F1174L for 48 hours and treated for 4 hours with ALK inhibitors, ceritinib (500 nM), alectinib (100 nM) and lorlatinib (100 nM) or an equivalent volume of DMSO. Western blots of pALK ^Tyr1604^ and α-tubulin were used to assess relative abundance in transfected HEK293 cells. **B, D, F.** pALK ^Tyr1604^ band intensity was quantified relative to α-tubulin from A, C and E. All graphs are representatives of two independent experiments (*n*=2).

**Supplementary Movie 1. Time-lapse imaging of EML4-ALK V3 WT droplet formation in HEK293 cells**

Time-lapse imaging of HEK293 cells transfected with YFP-EML4-ALK V3 WT. 25 z-sections of 1 μm step size were captured every second and time is shown in seconds.

**Supplementary Movie 2. Time-lapse imaging of EML4-ALK V3 kinase dead (KD) in HEK293 cells**

Time-lapse imaging of HEK293 cells transfected with YFP-EML4-ALK V3 D1270N. 25 z-sections of 1 μm step size were captured every second and time is shown in seconds.

**Supplementary Movie 3. Time-lapse imaging of EML4-ALK V3 WT in ceritinib-treated HEK293 cells**

Time-lapse imaging of HEK293 cells transfected with YFP-EML4-ALK V3 WT and treated with 500 nM ALK inhibitor, ceritinib. 25 z-sections of 1 μm step size were captured every second and time is shown in seconds.

**Supplementary Movie 4. Time-lapse imaging of EML4-ALK V3 WT in alectinib-treated HEK293 cells**

Time-lapse imaging of HEK293 cells transfected with YFP-EML4-ALK V3 WT and treated with 100 nM ALK inhibitor, alectinib. 25 z-sections of 1 μm step size were captured every second and time is shown in seconds.

**Supplementary Movie 5. Time-lapse imaging of EML4-ALK V3 WT in lorlatinib-treated HEK293 cells**

Time-lapse imaging of HEK293 cells transfected with YFP-EML4-ALK V3 WT and treated with 100 nM ALK inhibitor, lorlatinib. 25 z-sections of 1 μm step size were captured every second and time is shown in seconds.

**Supplementary Movie 6. Time-lapse imaging of EML4-ALK V3 WT by FRAP**

Time-lapse imaging of HEK293 cells transfected with YFP-EML4-ALK V3 WT and a ROI of 40×40 µm was photobleached at 100% argon 488 nm and 100% 405 nm laser power simultaneously. A second ROI of 40×40 µm was used as control (no photo-bleach) to represent the background.Single frames were captured every second and time is shown in seconds.

**Supplementary Movie 7. Time-lapse imaging of EML4-ALK V3 WT in alectinib-treated by FRAP**

Time-lapse imaging of HEK293 cells transfected with YFP-EML4-ALK V3 WT and treated with alectinib before imaging. A ROI of 40×40 µm was photobleached at 100% argon 488 nm and 100% 405 nm laser power simultaneously. A second ROI of 40×40 µm was used as control (no photo-bleach) to represent the background. Single frames were captured every second and time is shown in seconds.

